# Lipid-facilitated opening of the ADAM10 sheddase revealed by enhanced sampling simulations

**DOI:** 10.1101/2025.08.17.670722

**Authors:** Adrien Schahl, Nandan Haloi, Marta Carroni, Shengpan Zhang, Quentin James Sattentau, Erdinc Sezgin, Lucie Delemotte, Rebecca J Howard

## Abstract

ADAM10 is a crucial membrane-bound metalloprotease that regulates cellular physiology by cleaving and releasing membrane-anchored proteins, including adhesion molecules and growth factor precursors, thereby modulating cell signaling, adhesion, and migration. Despite its central role, its activation mechanisms are not fully understood. Here, we model how phosphatidylserine (PS) exposure during apoptosis triggers ADAM10 activation. We confirmed that PS externalization leads to shedding of CD43 from the surface of T cells via ADAM10 activity. Intriguingly, ADAM10 activation correlated with a loss of monoclonal ADAM10 antibody binding, suggesting a PS-induced conformational change that alters epitope accessibility. To explore this lipid-mediated conformational change of ADAM10, we employed molecular dynamics (MD) simulations to map the conformational landscape of ADAM10. Our simulations revealed that in the absence of PS, ADAM10 samples predominantly closed and intermediate states. By contrast, the presence of PS destabilizes the closed conformation, thereby favoring open states. We provide a mechanistic explanation for this PS-induced conformational change which drives ADAM10 activation and loss of mAb binding through conformational change. These findings offer new insights into the lipid-mediated regulation of ADAM10 and its conformational dynamics.

## Introduction

The “A Disintegrin And Metalloproteinase” (ADAM) family of membrane-anchored enzymes plays essential roles in many biological processes. One of their most important functions is to act as a cell surface sheddase, catalyzing the cleavage and release of multiple substrates including growth and differentiation factors, signaling molecules and cytokines. Dysregulated ADAM function is associated with a number of diseases including cancer, cardiovascular, neurodegenerative and inflammatory disorders^1^. Given the importance of the correct function of these enzymes, they are tightly regulated at transcriptional, translational and post-translational levels^2,3^. Post-translational regulation includes removal of an auto-inhibitory pro-domain during ADAM processing and complexation with a family of tetraspanins that modulate ADAM specificity and activity^4^. In the case of ADAM10 and 17, it has been recently proposed that phosphatidylserine (PS) flipping to the outer leaflet of the plasma membrane acts as a final trigger for activation^5,6^. PS is a phospholipid maintained primarily at the inner leaflet of the plasma membrane by flippases^7^. However, under conditions of cell activation and calcium flux, or cell death, PS is relocated to the outer leaflet by the action of scramblases^7^. Thus, scramblase activation likely indirectly modulates ADAM10/17 enzyme activity^8–11^. It has been hypothesized that the negative charge of PS interacts with a basic motif of amino acids in the Membrane-Proximal Domain of ADAM17 to activate the enzyme^12^. Mutation of this basic motif inhibits ADAM17 activation^12^, as does pharmacological inhibition of the interaction of PS with ADAM members^10,11,13^. Whilst this hypothesis is attractive, the structural modifications taking place in the ADAM enzymes leading to PS-mediated activation have yet to be defined.

Similarly to its close relative ADAM17, ADAM10 cleaves a wide variety of substrates with broad consequences for modulation of the immune system^14^. Of particular immunological importance, ADAM10 cleaves Notch, FAS-ligand, LAG3 and TIM3 in T cells, essential for T cell activation and function^14^. Amongst the more recently-discovered ADAM10 substrates are the transmembrane mucin-like molecules CD43, MUC-1 and CD162, which form a major component of the glycocalyx of immune cells including T cells^11^. Apoptotic cells must be rapidly cleared by phagocytes to prevent unwanted inflammation. Induction of apoptosis in T cells activates caspase-3 which in turn activates the scramblase XKR8 that flips PS to the outer leaflet^15^. This leads to the ADAM10-mediated shedding of mucins from the T cell surface^11^, reducing glycocalyx density and facilitating engulfment by phagocytes^11,16,17^. Given ADAM10’s role in immune modulation and tissue homeostasis, it is essential to reveal the molecular mechanisms by which PS exposure leads to ADAM10 activation.

Structurally, ADAM10 comprises seven subdomains: a pro-domain, a Metallo-Proteinase Domain (MpD), a Disintegrin-Like Domain (DD), a Cysteine-Rich Domain (CrD), a stalk domain (StD), a Transmembrane Domain (TmD), consisting of a single membrane-spanning helix, and an Intracellular Domain (IcD). The extra prodomain is cleaved upon maturation, only the six remaining subdomains are found in the plasma membrane (Fig. 1).

**Figure 1.**
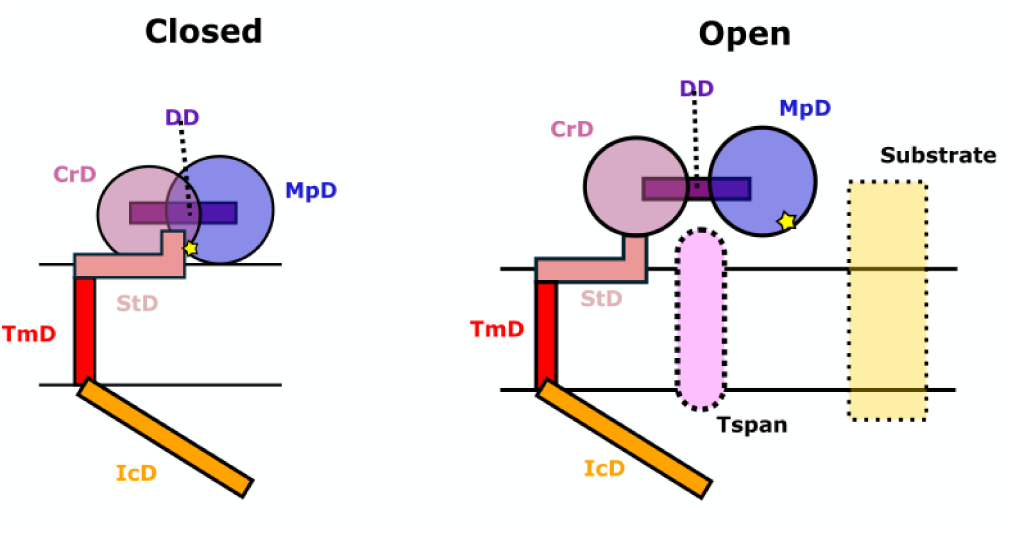
Subdomains of ADAM10, its interactors and two assumed conformations.

The MpD is responsible for ADAM10 enzymatic activity, and the CrD domains appear to be crucial for activation of ADAM10 and its closest homolog ADAM17^18^. In ADAM17, a small stalk section in the CrD called CANDIS (Conserved ADAM17 Dynamic Interaction Sequence) has been previously shown to form hydrophobic interactions with the membrane, thus triggering CrD structural changes leading to activation. Similarly, in ADAM10, the equivalent region of the protein (here termed StD) has been identified as important for its activation^6^. Particularly, a small patch of cationic residues, namely R657, K659 and K660, positioned in the StD, has been shown to interact with phosphatidylserine (PS), and ADAM10 activity was reduced upon mutation of these residues to asparagine.

To date, two distinct structures of ADAM10 extracellular domains have been resolved^19,20^. The first, obtained through crystallography, depicts a closed-like conformation where the StD occludes access to the MpD’s catalytic cavity. The second, revealed by cryo-EM, is a complex of ADAM10 with the scaffold protein tetraspanin (Tspan) and it shows a c-shaped open conformation exposing the ADAM10 catalytic cavity for substrate access. While this structural opening appears necessary for ADAM10 activity, the structural and dynamic details of this conformational transition, and its modulation by PS, remain to be elucidated.

Molecular dynamics simulations offer a powerful tool to bridge this knowledge gap by offering atomistic insights into dynamic molecular behaviors^21–23^. However, large-scale structural deformations, such as the opening of ADAM10, may require timescales on the order of tens of hundreds of microseconds to observe^24,25^. Enhanced sampling methods become instrumental in this scenario, enabling the efficient exploration of these conformational transitions^26^. For instance, adaptive sampling methods utilize swarms of short trajectories seeded to target specific transitions between states^27^. By employing Markov State Models (MSMs), these trajectories can be stitched together to construct a coherent and comprehensive view of the transition landscape^28,29^.

Here, we first experimentally confirmed that the presence of PS causes ADAM10 activation via imaging PS exposure on the cell surface and subsequent shedding of a fluorescently labelled ADAM10 substrate (CD43) during the apoptotic process. We also observed a basal activity of ADAM10 in healthy cells which, consistent with previous studies^11,30^, was reduced by ADAM10 inhibitors. Surprisingly, ADAM10 activation correlated with the loss of ADAM10 monoclonal antibody (mAb) clone 11G2 binding signal on the cell surface, suggesting a possible conformational change in ADAM10. Due to these observations, we hypothesized that ADAM10 assumes different conformational states with different levels of activity, and that PS engagement triggers a change towards more active states which no longer binds the antibody. To test these hypotheses, we performed MD simulations by leveraging the FAST (Fluctuation Amplification of Specific Traits) sampling method, yielding trajectories that could be analyzed using Markov State Modeling (MSM), thus producing free energy landscapes and transition rates between metastable states. This allowed us to characterize the energetic cost of ADAM10 conformational change (here termed “opening”) and to investigate how the presence of PS influences the energetics of this process. Our results reveal that, in the absence of PS lipids, ADAM10 samples predominantly closed and intermediate states and, to a small extent, the open state, suggesting that the opening is energetically accessible under these conditions. The presence of PS lipids in the outer membrane leaflet, on the other hand, completely destabilizes the closed states by capturing the CrD domain, thereby relatively stabilizing open conformations that promote ADAM10’s activity. Finally, we also show that the presence of PS brings the DD domain in close proximity to the membrane, which can account for the loss of mAb binding.

## Results and Discussion

### Apoptosis leads to loss of ADAM10 substrates and loss of ADAM10 antibody signal from the surface

To investigate the relationship between PS and ADAM10 activity, we fluorescently labelled ADAM10 and CD43 with antibodies on the human T cell leukemia CEM line. Apoptosis was induced using the kinase inhibitor staurosporine for different times and confirmed by antibody detection of active caspase-3 and externalized PS using fluorochrome-conjugated annexin-V. Analysis by confocal microscopy revealed that, as previously described^11^, fluorescence intensity of the mucin-like molecule CD43 reduced over time, but unexpectedly this was coordinate with an equivalent loss of ADAM10 labeling (Fig. 2A). The white arrow shows a rare early apoptotic cell at 1 h post-staurosporine treatment, which is positive for CD43, ADAM10 and PS. However, by 2 and 3 h post-staurosporine treatment all cells that were positive for Annexin-V and Caspase-3 were essentially negative for CD43 and ADAM10 labelling. This result was subsequently confirmed by flow cytometric analysis of activated Caspase-3 and ADAM10 co-labeling (Fig. 2B), revealing that >90% of cells positive for Caspase-3 were negative for ADAM10. We further quantified Caspase-3, ADAM10 and CD43 signal over time, and revealed a highly correlated and almost complete loss of CD43 and ADAM10 labelling by 4 h post-apoptosis induction that was negatively correlated with the kinetics of Caspase-3 activation (Fig. 2C).

**Figure 2.**
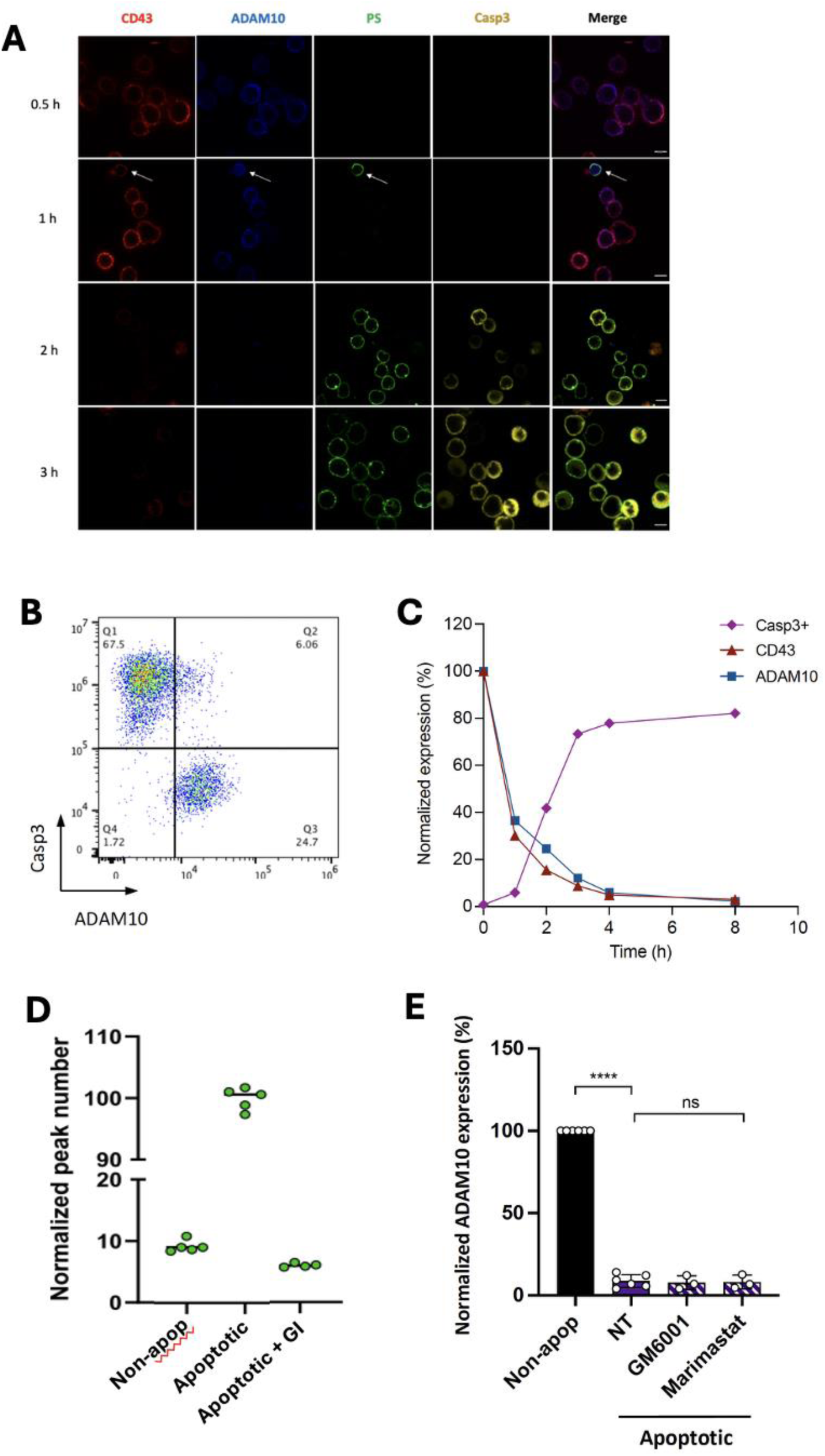
ADAM10 activity upon apoptosis induction. **A**) The fluorescence signal of anti-CD43 DFT1 antibody, anti-ADAM10 11G2 antibody, PS labelled with Annexin-V and Caspase-3 (Casp3) were recorded 0.5, 1, 2 and 3 h after apoptosis using fluorescence confocal microscopy. Scale bar = 7µm **B**) Casp3 vs ADAM10 signal analyzed by flow cytometry after 3 h apoptosis induction. **C**) Casp3, ADAM10 and CD43 signal over time quantified by flow cytometry expressed as % positive normalized to 100% at T = 0. **D**) Molecular quantification of shed CD43_Halo_ in the supernatant with single particle profiler normalized to apoptotic cells. **E**) ADAM10 signal quantified by flow cytometry in the absence or presence of broad-spectrum ADAM sheddase family inhibitors, n = 3 - 6 independent experiments expressed as % positive normalized to 100% ADAM10 expression on non-apoptotic cells.

We confirmed that loss of CD43 labeling was due to the shedding of the extracellular domain by quantifying shed CD43 extracellular domain from CEM stably transduced with CD43 fused to an N-terminal Halo tag (CEM-CD43_Halo_). CEM-CD43_Halo_ cells were labeled with fluorescent Halo substrate and induced to undergo apoptosis using staurosporine. CD43-Halo-ectodomain released into the supernatant after apoptosis was quantified using single particle profiler^31^ (Fig. 2D). At 3 h after apoptosis induction, we observed significantly higher numbers of CD43 soluble events compared to the healthy cell condition. Moreover, when we used a specific ADAM10 inhibitor (GI254023X, here termed GI), the number of soluble CD43 events was reduced dramatically to below the non-apoptosis condition (Fig. 2D).

Since the ADAM10-specific mAb was directly conjugated to a fluorochrome, the loss of signal over time cannot result from antigen-antibody internalization, as the mAb signal would have been maintained intracellularly and therefore still give positive signal by flow cytometry. ADAM10 has both autocatalytic activity and is targeted for cleavage by other ADAM proteases^32,33^. To exclude ADAM10 ectodomain cleavage as a mechanism of loss of antibody signal, we tested broad-spectrum metalloprotease inhibitors GM6001 and marimastat, which potently inhibit metalloprotease catalytic activity. Fig. 2E shows that these inhibitors had no effect on the loss of ADAM10 mAb binding during apoptosis, strongly suggesting that loss of mAb binding was due to conformational change in ADAM10 rather than its proteolytic cleavage.

These experiments yielded intriguing observations that necessitate further molecular understanding: i) how do PS (and other lipids) interact with ADAM10 to change its conformation? ii) how are the different conformations of ADAM10 linked to its function and antibody binding? In the following sections, we address these questions using MD simulations.

### Distribution of ADAM10 closed and open conformations obtained by enhanced sampling simulations

As we sought to understand the molecular mechanisms responsible for PS-mediated ADAM10 activation, we performed extensive MD simulations in different membrane environments. We hypothesized, considering available experimental structures, that ADAM10 activation would require the opening of the protein, accommodating both the Tspan scaffold protein and substrate binding. Such a conformational change is expected to take place over hundreds of µs to seconds, far beyond the accessible timescales of typical atomistic molecular dynamics simulations. We therefore decided to tackle this issue using enhanced sampling simulations, focusing specifically on the FAST adaptive sampling method. As detailed in Methods, this approach involves running successive sets of unrestrained MD simulations, iteratively using seeds selected based on expected progress towards a target state. The initial model was an experimental structure of ADAM10 in a so-called closed conformation (PDB ID: 6BE6), inaccessible to scaffold or substrate proteins, embedded in a homogenous 1-palmitoyl-2-oleoyl-glycero-3-phosphocholine (POPC) bilayer, thus mimicking the non-activated condition (Fig. 3A). We hypothesized that the transition from closed to open conformations, exemplified by the structure resolved in the presence of Tspan (PDB: 8ESV), could be modeled by using as target features the distances between the catalytic metalloproteinase domain (MpD) and the CrD/StD (Fig. 3A and Supp. Fig. S1). We thus defined this as a target for FAST sampling (see Methods) maximizing the sum of 127 pairwise Cα distances between these domains (Supp. Fig. S1, Supp. Tab. ST1). Over a total of 32 μs simulations, the summed interdomain features increased from ∼200 Å to ≥100 nm, convergence being reached after ∼8 generations (Fig. 3B).

**Figure 3.**
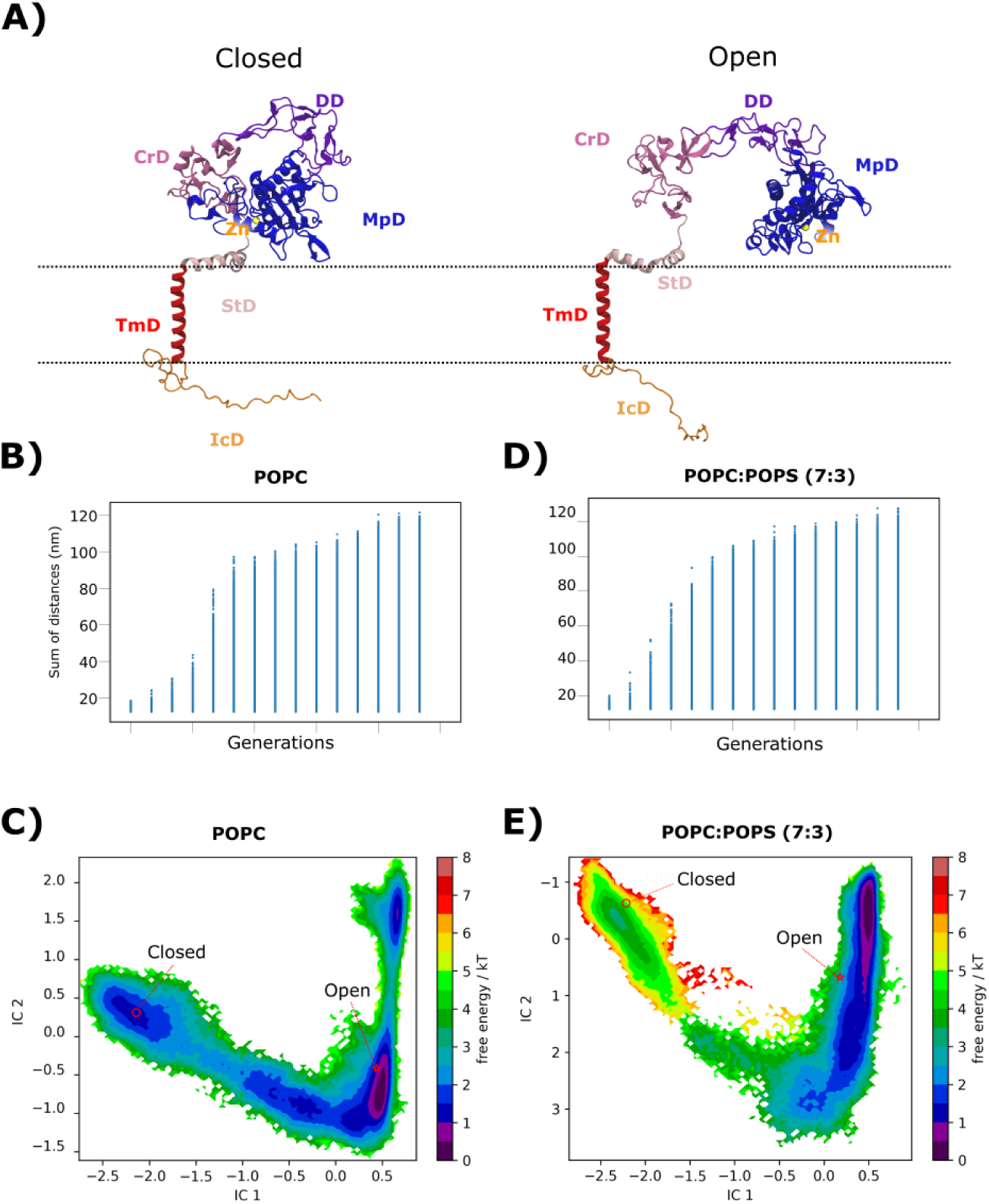
Distribution of ADAM10 closed and open conformations obtained by enhanced sampling simulations. **A)** Reference models of ADAM10 in an open (PDB ID: 8ESV, right) or closed (PDB ID: 6BE6, left) conformation. Coloring indicates the catalytic zinc ion (yellow) and the domains MpD (blue), DD (purple), CrD (magenta), StD (pink), TmD (red) and IcD (orange). Summed interdomain distances (Å) used to select seeds for successive generations of FAST sampling in the absence (**B**) and presence (**D**) of POPS. Free-energy landscapes built from MSMs of ADAM10 opening transitions in the absence (**C**) and presence (**E**) of POPS, plotted on the two slowest tICA components (IC2 vs. IC1), and colored by free energy according to the scale bar (*k*^-1^*T*^-1^). Reference structures shown in *A* in open (red star) and closed (red circle) conformations are projected onto each landscape.

We then performed time-lagged independent Component Analysis (tICA) based on interdomain-distance features. This analysis allowed us to characterize the slowest collective motions sampled by ADAM10. We then projected each simulation frame, along with reference structures, on the two-dimensional space formed by the two slowest motions (represented by the top two Independent Components IC1 and IC2) (Fig. 3C). The slowest component (IC1) appeared to capture substantial conformational changes between the initial closed structure and experimental open structure and was primarily characterized by the expansion between the MpD and the StD around the zinc-coordinated catalytic site. Transition from closed to open structures also involved displacement along the second slowest component (IC2), which appeared to capture variability between the MpD and CrD (Fig. 3C).

### Markov state models show POPS lipids favor opening transitions

We analyzed our trajectories using Markov State Modeling (MSM) and constructed a free-energy landscape for our sampled ADAM10 opening transition. Based on initial modeling using 200 microstates, the reference closed and open structures projected to distinct free-energy basins (Fig. 3C). Our central hypothesis is that the interaction of PS with ADAM10 changes the distribution of its conformations. To test this, we simulated the influence of POPS on ADAM10 opening. For this, we ran another series of MD simulations using the FAST-sampling protocol, with the protein embedded in a 7:3 mixture of POPC:POPS lipids, producing a further 32 μs of simulations. As for the POPC-only system, interdomain features converged to ≥100 nm within 8 generations (Fig. 3D). The slowest components (IC1, IC2) of the opening transition also appeared to capture similar structural features as in the POPC system. However, a tICA-based MSM revealed notable differences in the free-energy landscape of ADAM10, opening when embedded in the membrane containing POPS (Fig. 3E). In particular, the basin corresponding to the initial closed structure was disfavored relative to the one containing the experimental open structure.

To gain more insight into the thermodynamics and kinetics of this landscape, we then used the PCCA+ algorithm to build a 4-state coarse-grained MSM (Fig. 4A). Based on apparently progressive interdomain expansion, we termed these states Closed, Intermediate-closed (Ic), Intermediate-open (Io), and Expanded open (Eo, Fig. 4A). The initial experimental structure projected to the Closed state, while the open experimental structure projected to the Io state. Extracellular domains in the Ic state appeared to be partially expanded, while the Eo state exhibited even more extended domain interfaces than the experimental open structure. The Closed and Ic states were present in similar proportions, representing 20.0% and 20.6% respectively, the Io state was the most represented with 52.1% of the total distribution, while the Eo state accounted for the remaining 7.3%. This correlates well with our experimental results, where we showed that ADAM10 presents a basal activity (Fig. 2E). Mean first-passage times between states (MFPTs, the average time that takes to transition from one state to another) indicated that the Closed, Ic and Io states exchanged relatively rapidly, with ≤4.1 µs needed for a given transition; in contrast, transition times to the Eo state were on the order of 20 µs (Fig. 4B).

**Figure 4.**
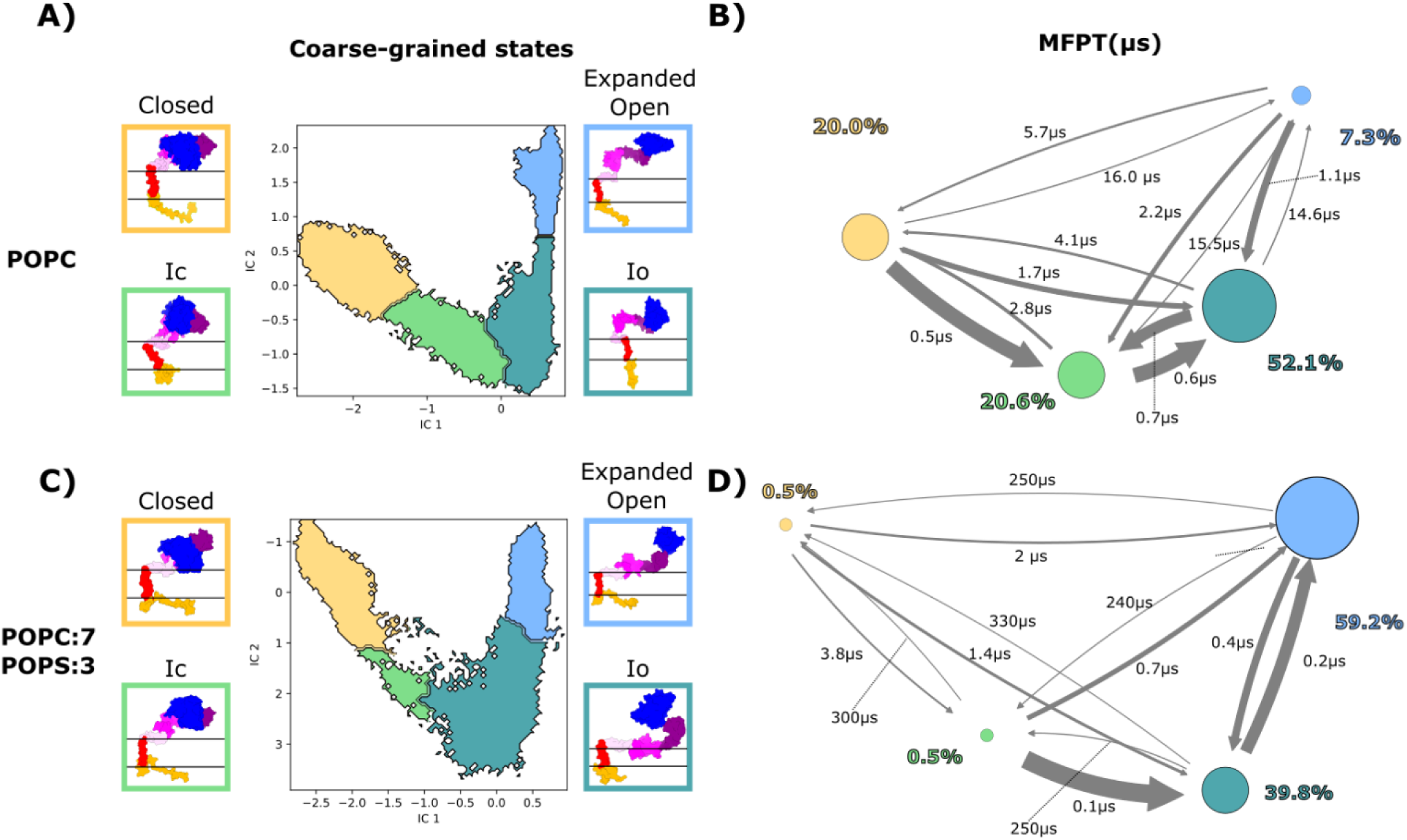
Markov state models show POPS lipids favor opening transitions. Coarse grained MSM computed with PCCA+ in **A)** POPC and **C)** POPC:POPS(7:3) conditions. Colored squares display a representative structure from the corresponding macro-state. Kinetic scheme representing exchanges between macro-states in **B)** POPC and **D)** POPC:POPS(7:3) conditions. The size of circle and arrows represents the relative population of each state and the mean first passage time (MFPT) between each state, respectively.

We then used the PCCA+ algorithm to build a 4-state model in the POPC:POPS conditions, characterized as Closed, Ic, Io and Eo (Fig. 4C). For this system, the Closed and Ic states represented only 0.5% each of the total distribution respectively, while the Io and Eo states accounted for 39.8% and 59.2% respectively (Fig. 4D). Transitions to the Io and Eo states were also accelerated in the presence of POPS, with MFPTs ≤2 µs, while transition times away from these states were estimated to be more than two orders of magnitude greater (>200 µs) (Fig. 4D). Finally, the Io and Eo states exchanged rapidly (< 0.5 µs).

### POPS limits CrD-membrane dissociation

To understand the molecular details of POPS-dependent sheddase opening, we next quantified contacts between membrane lipids and the membrane-proximal extracellular domains (MpD, StD and CrD) of ADAM10 in the different states. In the system with POPC alone, the CrD made substantial lipid interactions in both the Closed and Ic states, with a distribution centered around 10 direct contacts. However, the number of contacts with lipids was largely reduced in the Io and Eo states, suggesting that interactions of this subdomain with POPC are insufficient for this domain to remain membrane-anchored (Fig. 5A–C).

**Figure 5.**
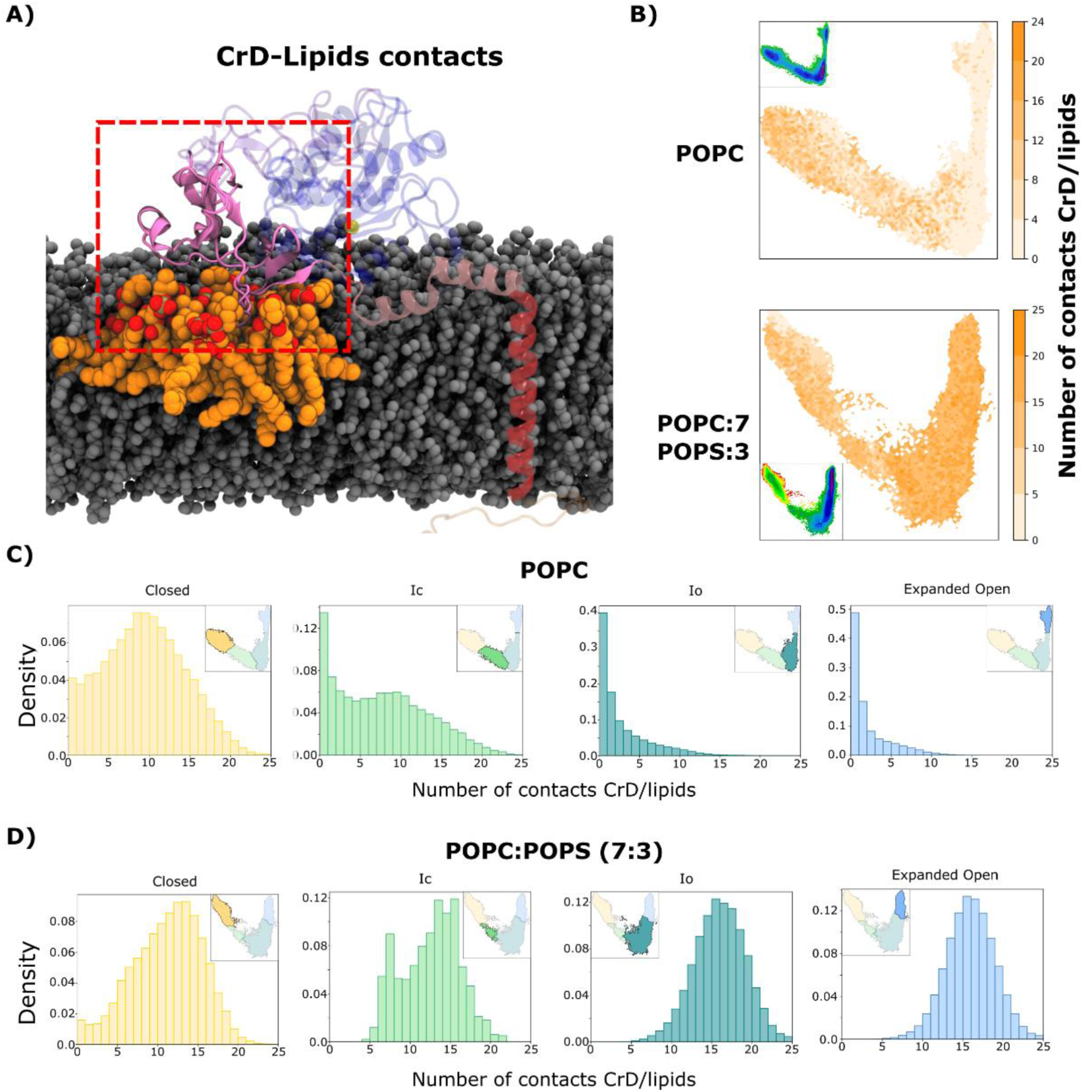
POPS limits CrD-membrane dissociation upon ADAM10 opening. **A)** Representation of lipids involved in contacts with the CrD. **B)** Projection of the number of lipids in contact with CrD over free energy maps in POPC and POPC-POPS simulations. Histogram of the number of contacts between lipids and CrD for each macro state obtained in in **C)** POPC and **D)** POPC:POPS(7:3) conditions.

In contrast, the CrD maintained ∼15 lipid interactions across all states in the presence of POPS (Fig. 5B, D). Thus, POPS charge interactions appeared to prevent dissociation of the CrD from the membrane during ADAM10 opening, consistent with facilitation of substrate-accessible states. As expected, all of this led to a lower position of the CrD with respect to the center of the membrane POPC:POPS(7:3), compared to the pure POPC condition (Supp. Fig. S2). A conserved cluster of basic residues (R657, K659, K660) within the StD (Supp. Fig. S3) had previously been implicated in promoting ADAM17 activity^12,34^. We evaluated the number of contacts between POPS lipids and these basic residues in ADAM10. The largest number of contacts is observed in the Closed state, suggesting that this catalytic patch helps to recruit POPS lipids around the CrD to anchor it to the membrane (Supp. Fig. S3). The MPD contacted the membrane in the Closed state but lost most lipid interactions in the other three states, both in the absence and presence of POPS, suggesting this domain is consistently solvated relatively early in the opening transition (Supplementary Fig. S4).

### POPS promotes accessibility to the Tspan scaffold and to the catalytic site

Having characterized a generalized opening transition of ADAM10 presumed to precede scaffold or substrate binding, we further investigated the accessibility of the functionally critical catalytic site across the same free-energy landscapes. We quantified catalytic-site accessibility on the basis of the center-of-mass distance between MpD residues 419–421 (surrounding the catalytic zinc ion) and StD residues 646–651 (Fig. 6A), largely represented by variations along IC1. Accessibility increased from ≤20 Å to 25–30 Å between Closed and Ic states in pure POPC, while staying around 20 Å in POPC:POPS conditions. In the absence of POPS, the Io state was characterized by a range of accessibility values centered around 65 Å, while the Eo state was relatively expanded to around 100 Å (Fig. 7B–C). In the presence of POPS, accessibility in the Open state was broadly distributed between 50 and 100 Å (Fig. 7B, D). Thus, the catalytic site appeared to be similarly accessible in the Io/Eo states in both systems, with the relative population of these two states increasing in the presence of POPS. The opening of ADAM10 is important to accommodate both substrate and Tspan binding. Thus, we decided to investigate distances between the two interfaces responsible for Tspan binding (Supplementary Figs. S1, S5). The evolution of these distances correlates well with the accessibility to the catalytic cavity.

**Figure 6.**
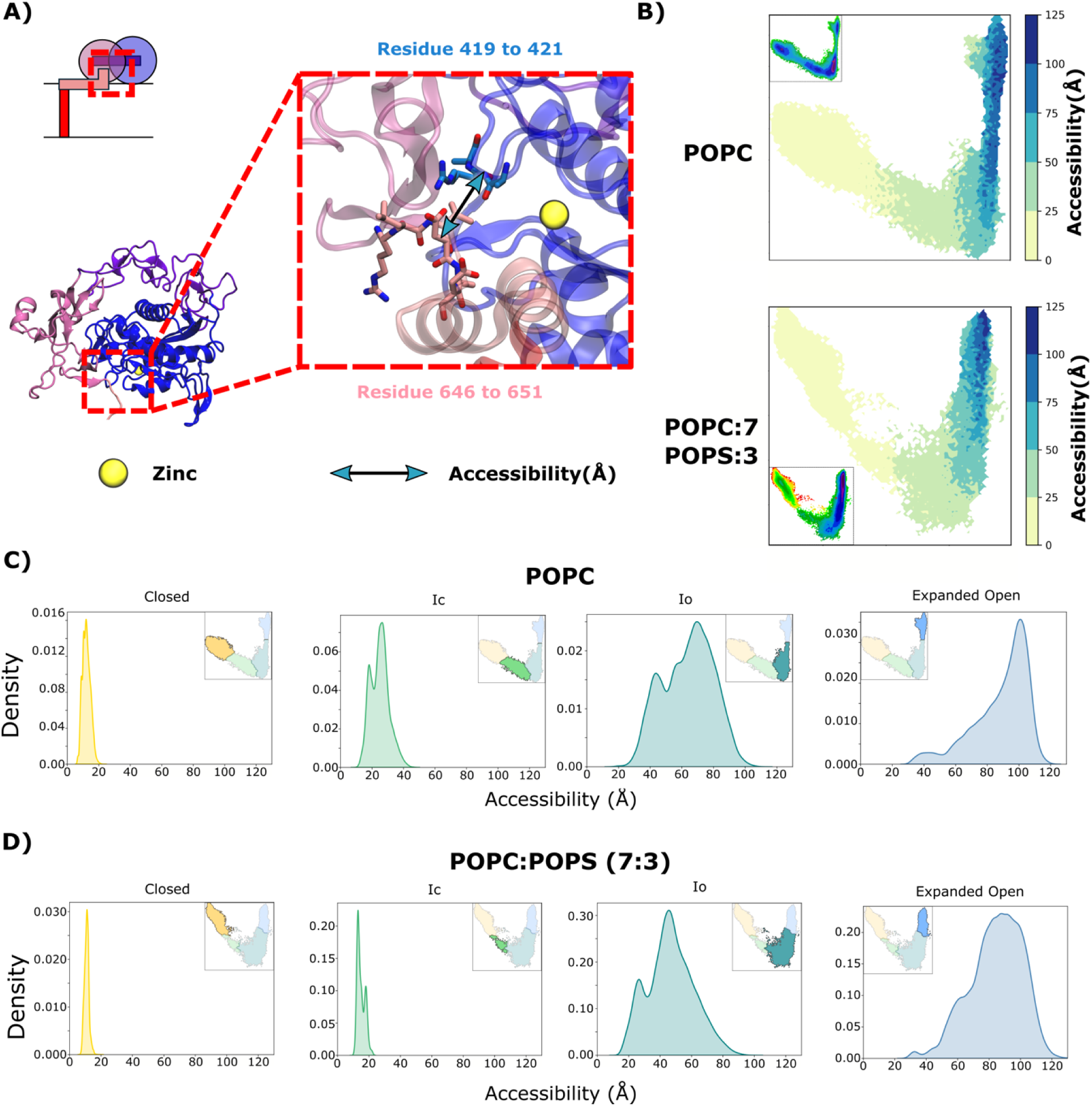
POPS promotes catalytic-site accessibility upon ADAM10 opening. **A)** Representation of the distance defining the accessibility of the catalytic cavity. **B)** Projection of this distance over free energy maps in POPC and POPC-POPS simulations. Distribution of this distance in each macro state obtained in **C)** POPC and **D)** POPC:POPS(7:3) conditions.

**Figure 7:**
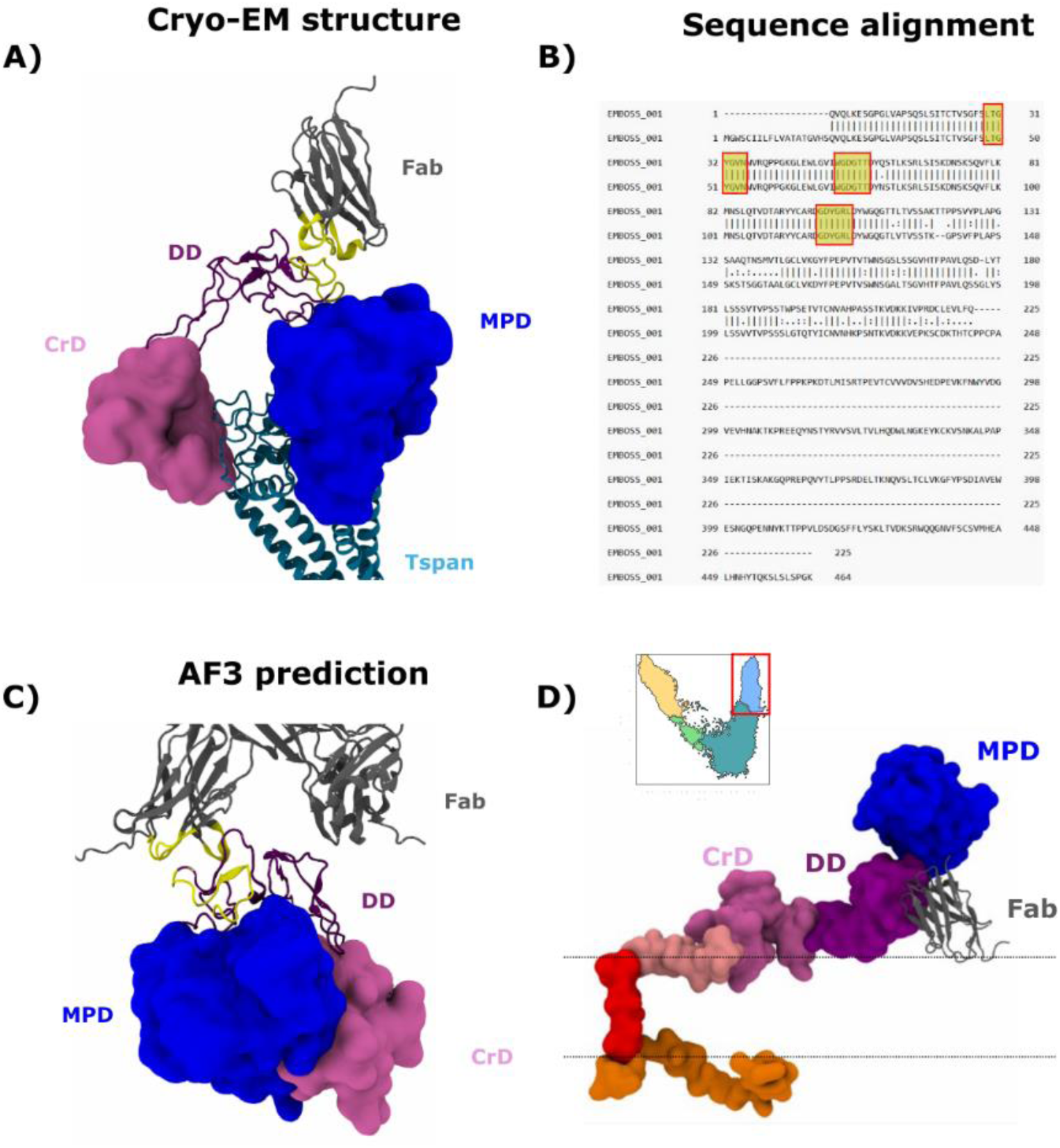
Conformational modulation of ADAM10 interaction with antibody. **A)** Representation of Fab_cryo_-ADAM10 binding in a cryo-EM structure (8ESV). **B)** Sequence alignment between the Fab used in our study and the one used in cryo-EM. Parts highlighted in yellow represent the antigen binding site. **C)** AF3 prediction of Fab_fluo_-ADAM10 binding. **D)** Projection of Fab binding on a representative structure extracted from the Open state, the most frequent one in POPC-POPS condition.

### POPS prevents antibody binding to ADAM10 by reducing DD-membrane distance

Having observed that ADAM10 changes conformation upon PS interaction, we tested whether this could influence the interactions of ADAM10 with the antibody used to detect it. To this end, we studied the binding interface between the DD and an antibody Fab fragment, as resolved in the cryo-EM structure used as a representative open state (PDB ID 8ESV) (Fig. 7A). We determined that the antigen binding site of our Fab heavy chain (Fab_fluo_) presented a sequence that was identical to the one used in the cryo-EM study (Fab_cryo_, Fig. 7B). We thus hypothesized that the Fab adopted a similar structure as Fab_cryo_ and that their antigen binding mode was similar. To test this hypothesis, we generated potential binding complexes structures using the AF3 server. Most of the predictions resulted in the Fab_fluo_ indeed displaying a similar binding site structure and binding close to the experimentally resolved binding site (Fig. 6C, Supp. Fig. S6). In the experimental structure, loops 48-54, 71-77 and 118-123 from the Fab_cryo_ (Fig 7A-B, highlighted in yellow), are bound to residues 475-498 from ADAM10. Particularly, residues 54-56 and residues 119-120 of the Fab_cryo_ form hydrogen bonds with respectively residues 498 and residues 491-492 from ADAM10. In the predicted structures, we observe the binding of loops 48-51, 71-77 from the Fab_fluo_ to residues 470-491 from ADAM10. In the predictions, we observed hydrogen bonds between residues 120 and 71,77 from the Fab and 470 and 493, respectively, from ADAM10. We then proceeded to superimpose the DD of the cryo-EM structure with 10 structures randomly extracted from the most populated state (Eo) in the POPC-POPS condition, to analyse how the Fab_cryo_ would be positioned with respect to the membrane. In 9 out of 10 extracted structures, the binding of Fab_cryo_ to ADAM10 led to clashes with the membrane (Fig. 7D, Supp. Fig. S7), suggesting that the geometrical changes caused in ADAM10 by PS lipids are responsible for the loss of antibody binding.

## Discussion

Structural and dynamical characterization of ADAM10 is crucial for understanding its function. Similar to other enzymes, its activity is closely linked to its propensity to adopt different conformational states and interconvert between them. Gaining insight into how ADAM10 transitions between different conformations can provide valuable information about its regulation and activation mechanisms, which are key to its biological roles.

Until recently, only a closed structure of ADAM10 had been defined^19^. However, a more recently published structure revealed an open and active conformation, enclosing a Tspan15 protein^20^. It thus appeared that the opening of ADAM10 was an important step towards its ability to cleave membrane substrates. As we and other groups previously showed that the presence of POPS in the outer membrane can enhance ADAM10 activity^6,11^, we hypothesized that the opening of ADAM10 would be modulated by the presence of these lipids in the plasma membrane. Our Markov State Models (MSM) analysis shows that, in a pure POPC membrane, the closed and open conformations are projected into two distinct metastable basins, with the open conformation appearing to be slightly more stable than the closed one. When analyzing the behavior of ADAM10 in a POPC:POPS (7:3) membrane mixture, we observed striking differences in its conformational dynamics compared to a pure POPC membrane. Most notably, the closed structural basin was severely destabilized relative to the open one. These results suggest that enhancing the opening of ADAM10 promotes its activity.

To quantify this process, we further constructed a coarse-grained MSM comprising four states, ranging from the most closed to the most open: Closed, Ic, Io, and Eo. Interestingly, while the experimental closed structure aligns with the Closed state, the experimental open structure corresponds to the Io state. This suggests that ADAM10 may adopt even more extended conformations in solution. Notably, the Io state accounts for 52% of the structures, making it the most prevalent conformation in pure POPC. This observation sheds new light on ADAM10 structure and dynamics, as it was thought to be mainly in closed state at the surface of the cell^19,35^, potentially in dimer form. After obtaining the closed structure, Seegar and co-authors suggested that ADAM10 could present a transient opening leading to its activation. Here, our models support the concept that ADAM10 can indeed adopt an open conformation in the membrane, without the need for an external stimulus. Our data also confirm previously reported basal activity of ADAM10 that can be reduced by the addition of an inhibitor.

Another dimension of ADAM10’s regulation regards its interactions with Tspan proteins, as observed biochemically^36^ and in crystallographic^37^ and cryo-EM structures^20^. Proteins from the Tspan family are known regulators of the ADAM10 protein, especially regarding its surface expression^38^. In this context, Tspan15 upregulates the expression of ADAM10 and limits its endocytosis^36^. Nonetheless, the question of the process of assembly between ADAM10 and

Tspan proteins remains an open question, especially since a closed, autoinhibited structure of ADAM10 has been crystallized. Our results confirm that ADAM10 encloses Tspan and that the formation of the complex may occur without any interaction with PS lipids, e.g. in the healthy plasma membrane. Here we note that the percentage of open structures (50%) in our model does not explain quantitatively the reduction of activity that we measure in our experiments. As Tspans modulate the selectivity of cleavage of ADAM10, a possible explanation would then lie in the competition between associations of the protein with different Tspans, as has been shown previously^38,39^. The percentage of substrate cleaved in our experiment would then correspond to a subset of the open structures that we observe in our simulations. In this context, the fast switch between open and closed states allows ADAM10 to present a basal activity that would be linked to its ability to bind Tspan and/or to access the substrate (Fig. 8).

**Figure 8.**
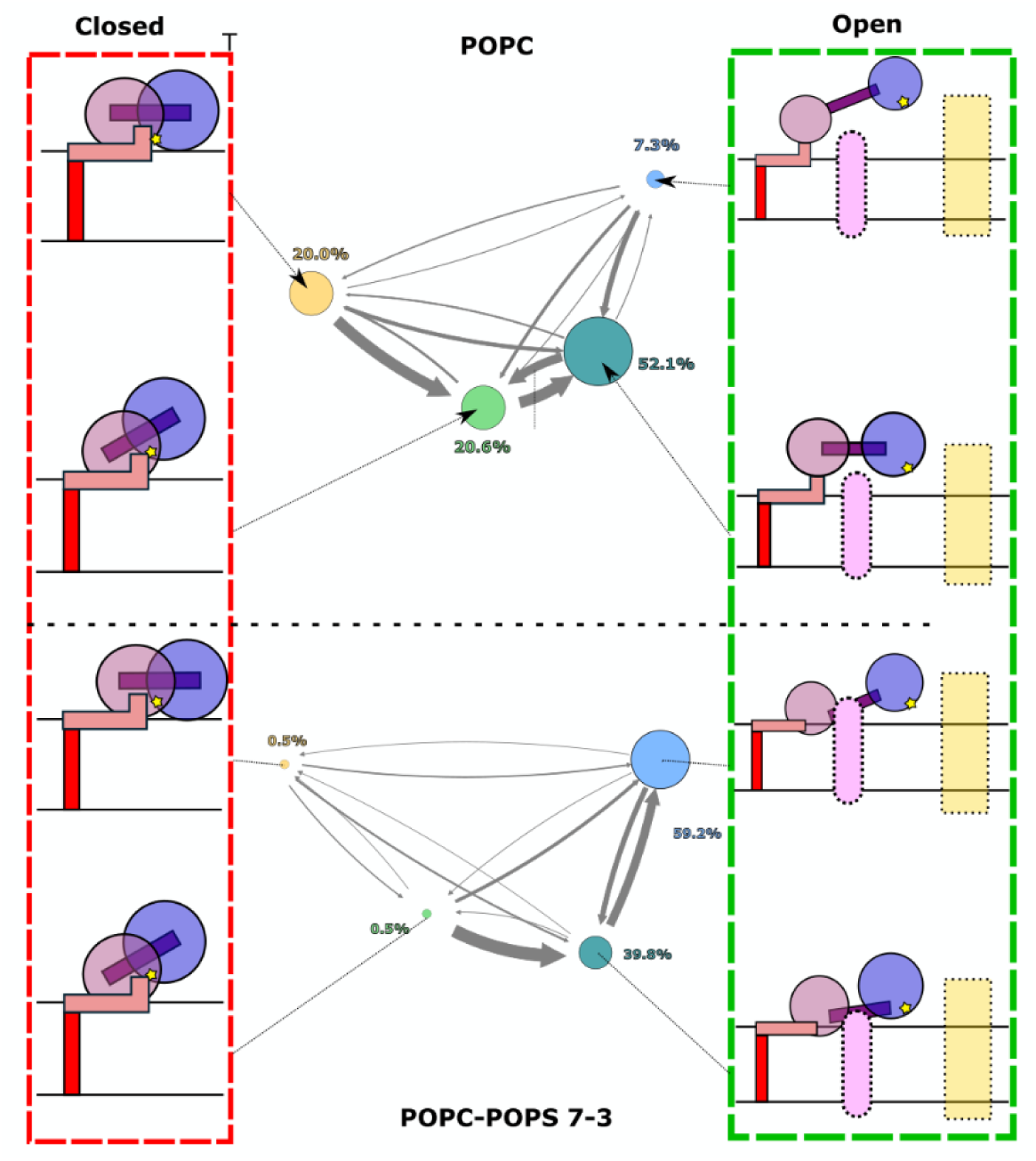
Schematic of conformational transitions in ADAM10 opening in the absence or presence of POPS. Without POPS, ADAM10 will oscillate between closed and open states in the membrane, with a similar proportion of each structure in each state. In the closed states, the catalytic site (yellow star) is not accessible to any substrate, and the binding to proteins of the Tspan family will be unlikely. Upon addition of POPS, closed states will disappear, allowing ADAM10 to bind Tspan (pink oval) or to cleave divert substrates(yellow rectangle).

Upon the addition of PS lipids in our model, we observed a drastic change in the percentage of open structures, with the Io and Eo states accounting now for 99% of the observed structures. Whilst the autoinhibitory forms of ADAM10 become non-existent, the enzyme activity increases in relation to the increasing presence of open forms that can either bind Tspans or cleave diverse substrates. In their work, Seegar and co-authors studied the binding interface of ADAM10 with clone 8C7 antibody, which has an enhancing effect on ADAM10 activity. They demonstrated that in the autoinhibitory form, the MPD of ADAM10 would clash with the heavy chain of 8C7, explaining why this latest enhances the activity of the sheddase. This, added to our results, further highlights the correlation between the need of opening of ADAM10 and its ability to exert its function, highlighting the tight relation between its structure and activity.

The key factor in ADAM10 modulation by PS lipids is their interaction with the CrD. These persistent interactions with the membrane could explain why it is difficult for the system to revert from the Io state to the Ic state, as the CrD remains anchored and restricts conformational reversibility. This ADAM10 behavior appears comparable with that of ADAM17, the activity of which was modulated by changes of CrD conformation mediated by interactions with negatively charged lipids^21,41,42^, which suggests a common feature of this family of proteins. Surprisingly, the previously characterized cationic patch present in ADAM17 (R657, K659, K660) displayed increased interactions with POPS lipids specifically in the Closed state. This suggests that these residues may also help recruit lipids to ADAM10, which leads to the anchoring of the CrD domain and the potential stabilization of its association with the membrane. The PS-mediated repositioning of the CrD also influences the position of the DD, bringing it closer to the membrane in the open state. In this state, the binding of 11G2 FAb would become impossible, due to steric clashes with the membrane, thus explaining the loss of ADAM10 signal that we observed in our experiments.

Despite these positional differences, the opening of ADAM10, characterized by the distance between the StD and the catalytic cavity (Fig. 6), remained consistent with the POPC-only simulations. This indicates that the four states observed in the POPC: POPS membrane retain a similar structural framework to those identified in pure POPC conditions. Together, these findings highlight that the presence of POPS not only shifts the equilibrium towards open conformations but also influences specific membrane/protein interactions that are likely crucial for the activation and function of ADAM10.

In conclusion, the addition of POPS shifts ADAM10 toward more open conformations, coordinate with a predicted enhancement of its activity. The closer proximity of the CrD and MpD to the membrane in this condition facilitates interactions with tetraspanins, supporting the hypothesis that POPS promotes ADAM10’s readiness to enclose Tspan proteins. This highlights a dual role for POPS in modulating both its activity and structural accessibility. Understanding how ADAM10 conformational states influence catalytic activity and specificity will have utility in designing agents to either enhance or inhibit enzyme activity, with direct therapeutic relevance.

## Material and methods

### Flow cytometric analysis of antibody binding to ADAM10 on healthy and apoptotic T cells

Acute T cell lymphoid leukemia (T-ALL) line CEM was obtained from the Sir William Dunn School of Pathology cell bank. CEM was grown in complete RPMI (ThermoFisher 11875093) containing 10% FCS (Sigma-Aldrich F9665), 1% penicillin/streptomycin (ThermoFisher 15140122) at 37 °C/5% CO_2_. CEM were counted and resuspended at a concentration of 2–3 × 10^6^ cells/mL in complete RPMI. Staurosporine (Cambridge Bioscience S-7600) was pre-titrated to 10 μM giving ∼50% apoptosis at 3 h post-treatment. Cells were washed with PBS and collected for further manipulation. Cells were centrifuged for 5 min at 400 x g. Pellets were resuspended in 100 μL cold annexin-V binding buffer (BD Pharmingen 556454) for 20 min at 4 °C in the dark with annexin V-FITC (Biolegend 640906) used at 1:100, near-IR fixable viability dye (Invitrogen L10119) at 1:1000, fluorophore-conjugated anti-human CD43 sialic acid-independent clone L10-APC (Invitrogen 10746863) used at 2 μg/mL and anti-human ADAM10-BV421 clone 11G2 (BD Biosciences 742787) used at 2µg/mL. All labelling was done in conjunction with the corresponding concentration-matched isotype control antibody. After labeling, cells were washed with cold annexin V binding buffer where appropriate, centrifuged for 2 min at 400 x g at 4 °C, and fixed with 4% paraformaldehyde (Sigma-Aldrich 158127) for 10 min at RT. Fixed cells were washed with PBS, permeabilized with perm buffer (Biolegend 421002), and labeled with anti-human active caspase-3 antibody clone Asp175 (Cell Signaling Technologies 9661) used at 1:400. After 30min incubation at 4 °C, cells were washed with FACS wash buffer (PBS, 2% FCS) and incubated with anti-Rabbit IgG (H+ L) Alexa Fluor 546 (Invitrogen, A11010) used at 4 μg/mL (1:500) where required for 30 min at 4 °C. After washing, cells were analyzed by flow cytometry using a Cytoflex LX flow cytometer (Beckman Coulter) and data processed using the FlowJo-V10 software (FlowJo, LLC). Isotype controls and Fluorescence Minus One (FMO) controls were performed for all colours to gate on positive and negative populations. Gating on the relevant cell population was set according to Forward Scatter (FSC) and Side Scatter (SSC) before doublet and near-IR fixable viability dye (Thermo Fisher) to allow exclusion of dead cells with dye-permeable membranes. Subsequent gating for all samples was carried out on annexin V and/or activated caspase-3 labeling to differentiate apoptotic from non-apoptotic cells within the same total cell population. Broad spectrum metalloprotease-specific inhibitors (GM6001, R&D Systems 2983; Marimastat, R&D Systems 2631) were each added at the pre-determined optimum concentration of 10 μM for 40 min prior to the addition of staurosporine, and maintained during the experiment.

### Confocal microscopy of healthy and apoptotic T cells

CEM were counted and apoptosis induced as for flow cytometric analysis. Cells were washed with PBS resuspended in 100 μL cold annexin-V binding buffer (BD Pharmingen 556454) for 20 min at 4 °C in the dark with annexin V-FITC (Biolegend 640906) used at 1:100, Alexa Fluor 647 conjugated anti-human CD43 sialic acid-dependent clone DFT1 (Santa Cruz Biotechnology, sc-6256) used at 2 μg/mL and anti-human ADAM10-BV421 clone 11G2 (BD Biosciences 742787) used at 2 μg/mL. After labeling, cells were washed with cold annexin V binding buffer where appropriate, centrifuged for 2 min at 400 x g at 4 °C, and fixed with 4% paraformaldehyde (Sigma-Aldrich 158127) for 10 min at RT. Fixed cells were washed with PBS, permeabilized with perm buffer (Biolegend 421002), incubated with blocking buffer (5% BSA, 0.015% Triton in PBS) for 45 min and labeled with anti-human active caspase-3 antibody clone Asp175 (Cell Signaling Technologies 9661) used at 1:400. After 60min incubation at 4 °C, cells were washed with PBS and incubated with anti-Rabbit IgG (H+ L) Alexa Fluor 546, (Invitrogen, A11010) used at 4 μg/mL (1:500) for 30 min at 4 °C. Cells were washed with PBS and mounted in 5 μL of DAKO mounting medium, covered with coverslips, dried overnight and sealed with nail polish. Images were acquired on a Zeiss 880 Airyscan confocal microscope in superresolution mode, and analyzed using ImageJ software.

### Analysis of CD43 shedding by ADAM10

CEM-CD43_Halo_ cells were prepared as previously described^11^. Cells in logarithmic growth phase were washed twice and resuspended in fully supplemented RPMI medium (without antibiotics). All washing and centrifugation steps were performed at 1500 rpm for 1 min. Cells were labeled with 0.33 µM HaloTag® Alexa Fluor® 488 Ligand (Promega) to stain the CD43 ectodomain, gently mixed, and incubated at 37 °C for 1 h. After labeling, cells were washed twice and resuspended in 500 µl of unsupplemented, phenol red-free L15 medium (Thermo Fisher). Cells were either left untreated, treated with 10 µM staurosporine at 37 °C for 3 h, or pre-treated with GI inhibitor (1:1000 dilution) at 37 °C for 45 mins followed by 10 µM staurosporine for 3 h. The total incubation time was kept constant across all conditions. Cells were pelleted, and the supernatant was collected, centrifuged again, and transferred to a 0.5 ml 40K MWCO Zeba™ spin desalting column (Thermo Fisher), pre-washed twice with PBS. The flowthrough was clarified twice more using 40K columns, and the final flowthrough was used for fluorescence correlation spectroscopy (FCS) measurements. Peaks were counted using FCS measurements which were performed using a Zeiss LSM 980 confocal microscope. Excitation of Alexa Fluor 488 was achieved using a 488 nm argon ion laser. A 40×/1.2 NA water immersion objective was used for focusing. For each sample, 10 fluorescence intensity curves (10 s each) were recorded. Laser power was set to 1% of the total output, corresponding to approximately 10 µW. Intensity traces were analyzed to count fluorescence peaks using single particle profiler software^31^.

### MD simulations system preparation

As a starting point for the simulations, we used the x-ray structure of ADAM10 (PDB:6BE6)^19^. The missing residues (670-748) were reconstructed using Alphafold2^43,44^, resulting in a single transmembrane helix with 2 juxtamembrane helices from residues 654 to 677, that we term StD. The initial systems were built using CHARMM-GUI^44^. The final ADAM10 model was embedded in a lipid bilayer composed either of only POPC or a mixture of POPC:POPS(7:3). The system was then solvated in a box of explicit water molecules and neutralized in 150 mM KCl. To increase the sampling of POPC-POPS systems, we generated 20 different initial membrane repartitions by using the membrane mixer plugin in VMD. The system compositions are described in Table 1:

**Table 1:** Composition of simulated systems.

| System | Lipids | Water | Number of atoms | duration |
| --- | --- | --- | --- | --- |
| POPC | 761 | 150k | 550k | 32μs |
| POPC:POPS(7:3) | 560:240 | 150k | 550k | 32μs |

MD simulations of ADAM10 were performed using GROMACS 2023.2^45^. The CHARMM36^46,47^ force field was employed in combination with the TIP3P^48^water model. The systems underwent energy minimization using the steepest descent algorithm until the energy gradient converged to a threshold of 0.01 kcal/mol/Å. The solvent and membrane were then allowed to relax in the NVT ensemble at 300K for 625 ps. The system then underwent NPT equilibration during 3 ns, first increasing the timestep from 1 fs to 2 fs, then progressively removing all constraints applied on protein and lipids. The LINCS algorithm^49^ was used for bond constraints.

During equilibration, the Berendsen thermostat^50^ and barostat (when applicable) were used. For production, the v-rescale thermostat^51^ and c-rescale barostat^52^ were used for temperature and pressure control, respectively. For equilibration as well as production runs, a cutoff distance for non-bonded interactions was set at 1.2 nm, utilizing a cut-off van der Waals (vdW) type with a force-switch modifier. The switching distance was configured to 1.0 nm.The Coulombic interactions were calculated using the Particle Mesh Ewald (PME) method^53^, with a cutoff distance of 1.2 nm.

### Adaptive sampling

We performed adaptive sampling based on the Fluctuation Amplification of Specific Traits (FAST) method^27^ to enhance the exploration of the conformational landscape. Briefly, the method consists of setting up a swarm of short simulations, clustering structures among the resulting trajectories, and selecting new structures among them to set up a new swarm. The new structures were chosen based on a reward function defined as the sum of 127 pairwise distances spread among 4 interfaces between the catalytic domain of ADAM10 and the other subdomains (Supp. Fig. 1, Supp. Tab. ST1). We then generated 16 generations of 20 simulations, each with a length of 50 ns. Once the overall sampling done, simulations from all seeds were extended to ensure a better sampling, amounting to 32 µs of sampling for each system.

### Markov State Models

To analyze the simulation data, Markov State Models (MSMs) were constructed using pyEmma^54^. To this aim, we performed Time-lagged Independent Component Analysis (TICA)^55^ on the set of pairwise distances described above, and added pair-wise distances corresponding to distances between the regions of ADAM10 interacting with Tspan15 (Supp. Fig. 1, Supp. Tab. ST1). As previously described^24,56^, faster dimensions represented by a single Gaussian distribution were discarded, and we decided to focus on two independent components for the analysis. After VAMP-2 validation^57^, 200 microstates were clustered using k-means^58^ clustering(Supp. Fig. 9). We used bayesian MSM^59^, on the implied timescales on a range of lag times going from 0.1 to 50 ns, to validate the use of a 10 ns lag time^60^(Supp. Fig. 10). Then, we performed PCCA+ analysis^61^ to build a 4 and 3 states Coarse-grained MSM model of the POPC and POPC-POPS systems respectively. This latest model was validated using the Chapman-Kolmogorov test^60^ (Supp. Fig. 11). Relevant observables were computed using VMD^62^, and results were displayed using seaborn^63^ and matplotlib^64^.

## Data availability

All data will be available upon publication at FigShare.

## Acknowledgements

Authors acknowledge SciLifeLab Research Environment Development (RED) grants. We thank the SciLifeLab Advanced Light Microscopy facility and National Microscopy Infrastructure (VR-RFI 2016-00968) for their support on imaging. ES has been supported by Swedish Research Council Grants (grant no. 2020-02682, 2024-02993 and 2024-00289), Wellcome Leap’s Dynamic Resilience Program (jointly funded by Temasek Trust), Karolinska Institutet (2024-03250; 2024-03341; 2022-00803; 2020-00997), Cancer Research KI (2024-03488), Human Frontier Science Program (RGP0025/2022), Longevity Impetus Grant from Norn Group, Hevolution Foundation and Rosenkranz Foundation. LD was supported by the Knut and Alice Wallenberg Foundation and Swedish Research Council grants VR 2019-02433 and 2022-04305.

MD simulations were performed using computing facilities of the Karolina, LUMI and Discoverer Supercomputer through EuroHPC (grant nos. EHPC-REG-2023R01-103, EHPC-REG-2022R03-223 and EHPC-REG-2022R03-219, respectively) and the Swedish National Infrastructure for Computing (SNIC 2022/3-40, NAISS 2024/5-71, NAISS 2024/3-47, NAISS 2025/1-34) and supported by BioExcel (EuroHPC grant no. 101093290). N.H. was supported by a Marie Sklodowska-Curie Postdoctoral Fellowship (grant no. 101107036). Q.J.S was supported by an award from the UKRI MRC, and SZ by a studentship from the China Scholarship Council 0 University of Oxford. Confocal microscopy and flow cytometry were carried out at the Sir William Dunn School, University of Oxford core facilities with kind expert assistance from Robert Hedley, Errin Johnson and Alan Wainman.

