## Supplementary Figures and Tables for "Lipid-facilitated opening of the ADAM10 sheddase revealed by enhanced sampling simulations"

### Supplementary Information for "Lipid-facilitated opening of the ADAM10 sheddase revealed by enhanced sampling simulations"

Adrien Schahl[1,*], Nandan Haloi[1], Marta Carroni[2], Shengpan Zhang[3,4], Quentin James Sattentau[4,5], Erdinc Sezgin[6,*] , Lucie Delemotte[1,*], Rebecca J Howard[2,*]

[1] Science for Life Laboratory, Department of Applied Physics, KTH Royal Institute of Technology, 12121 Solna, Stockholm, Stockholm County 11428, Sweden

[2] Science for Life Laboratory, Department of Biochemistry and Biophysics, Stockholm University, Solna, SE-171 65, Sweden

[3] The Kennedy Institute of Rheumatology, University of Oxford, Roosevelt Drive, Oxford OX3 7FY, UK.

[4] Sir William Dunn School of Pathology, University of Oxford, Oxford, OX1 3RE, UK.

[5] The Max Delbrück Centre for Molecular Medicine, Campus Berlin-Buch, 13125 Berlin, Germany.

[6] Science for Life Laboratory, Department of Women's and Children's Health, Karolinska Institutet, Tomtebodavägen 23, 17165 Solna, Sweden

*** Correspondence**

*Additional validation SI figures to be referenced in Methods…*

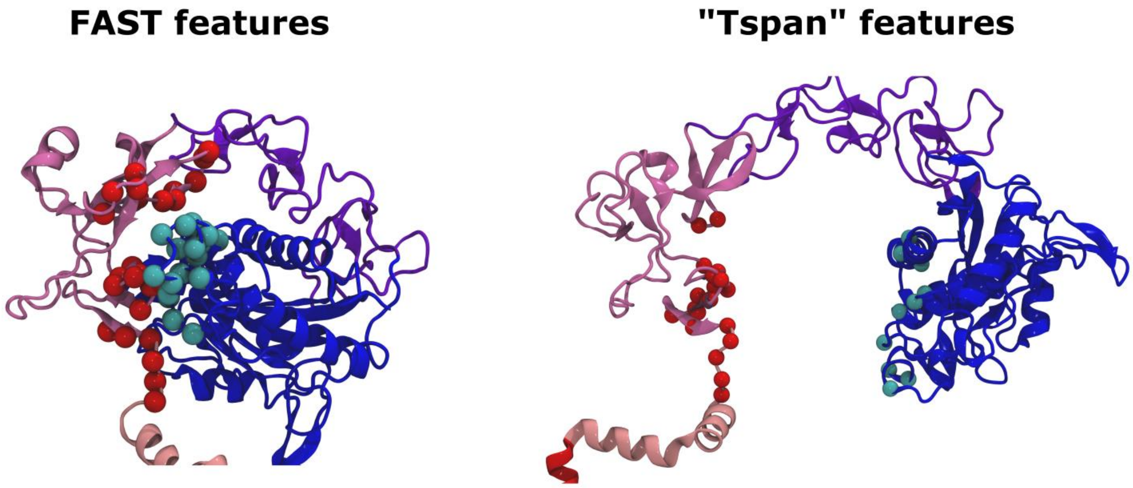


**Supplementary Figure S1. Features used for adaptive sampling and MSM building.** Left : Residues of the MpD (Cyan), CrD/StD (Red), used to define interdomain distances to select seeds for successive generations of FAST sampling. Right:  Residues of the MpD (Cyan), CrD/StD (Red), added as features for the construction of MSM. The numerotation of residues is displayed in Supplementary Table ST1.

**Supplementary Table ST1. Features numerotation used for adaptive sampling and MSM building** Numerotation of residues involved in the definition of pairwise distances, colored according to Supplementary Figure S1

| Features | CrD/StD residues number | MPD residues number |
|---|---|---|
| FAST sampling | 564-570 | 374-376; 424-426 |
| | 601-604 | 407-412 |
| | 630-636 | 401-405; 413-414 |
| | 646-651 | 419-421 |
| Tspan chelation | 556-557;<br>627-631; 635;<br>638-642; 646;<br>648-652 | 248-249; 251;<br>375; 378;<br>407; 410-411;<br>425; 430 |

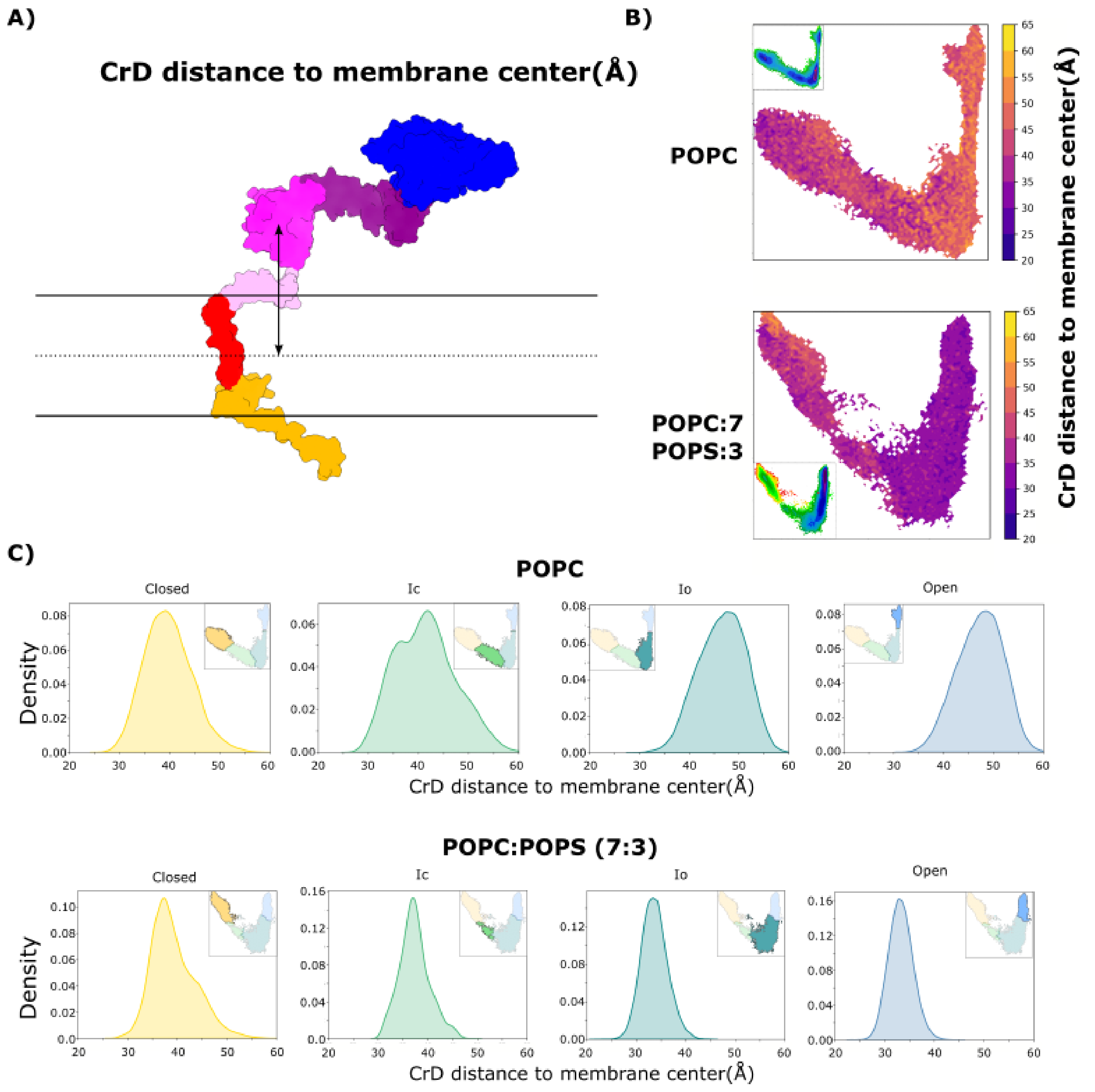


**Supplementary Figure S2. The CrD stays closer from membrane lipids in presence of POPS. A)** Representation of the distance between the CrD and the center of the membrane **B)** Projection of this distance over free energy maps in POPC and POPC-POPS simulations. **C)** Distribution of this distance in each macro state obtained in both conditions.

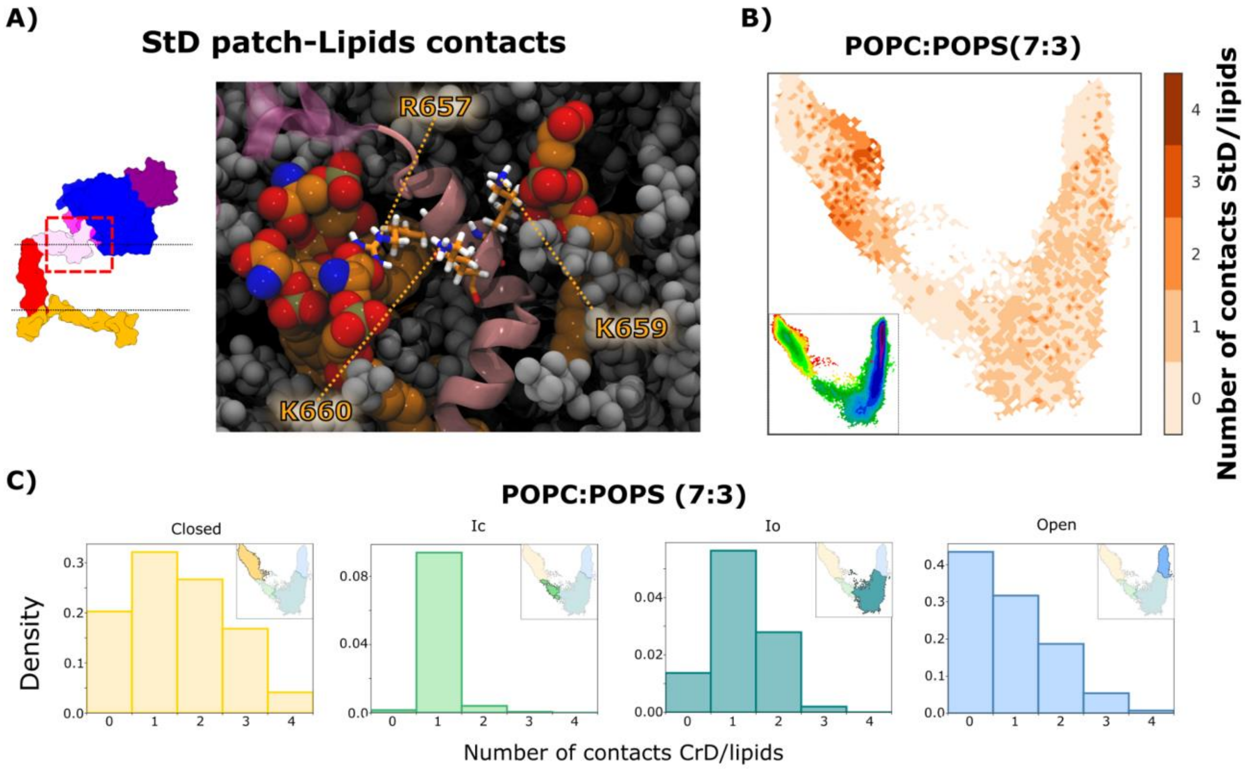


**Supplementary Figure S3. POPS interactions with basic residues in the ADAM10 StD. A)** Representation of the contacts between POPS lipids and basic residues of the StD **B)** Projection of the number of contacts over free energy maps in POPC-POPS simulations. **C)** Distribution of the number of contacts in each macro state.

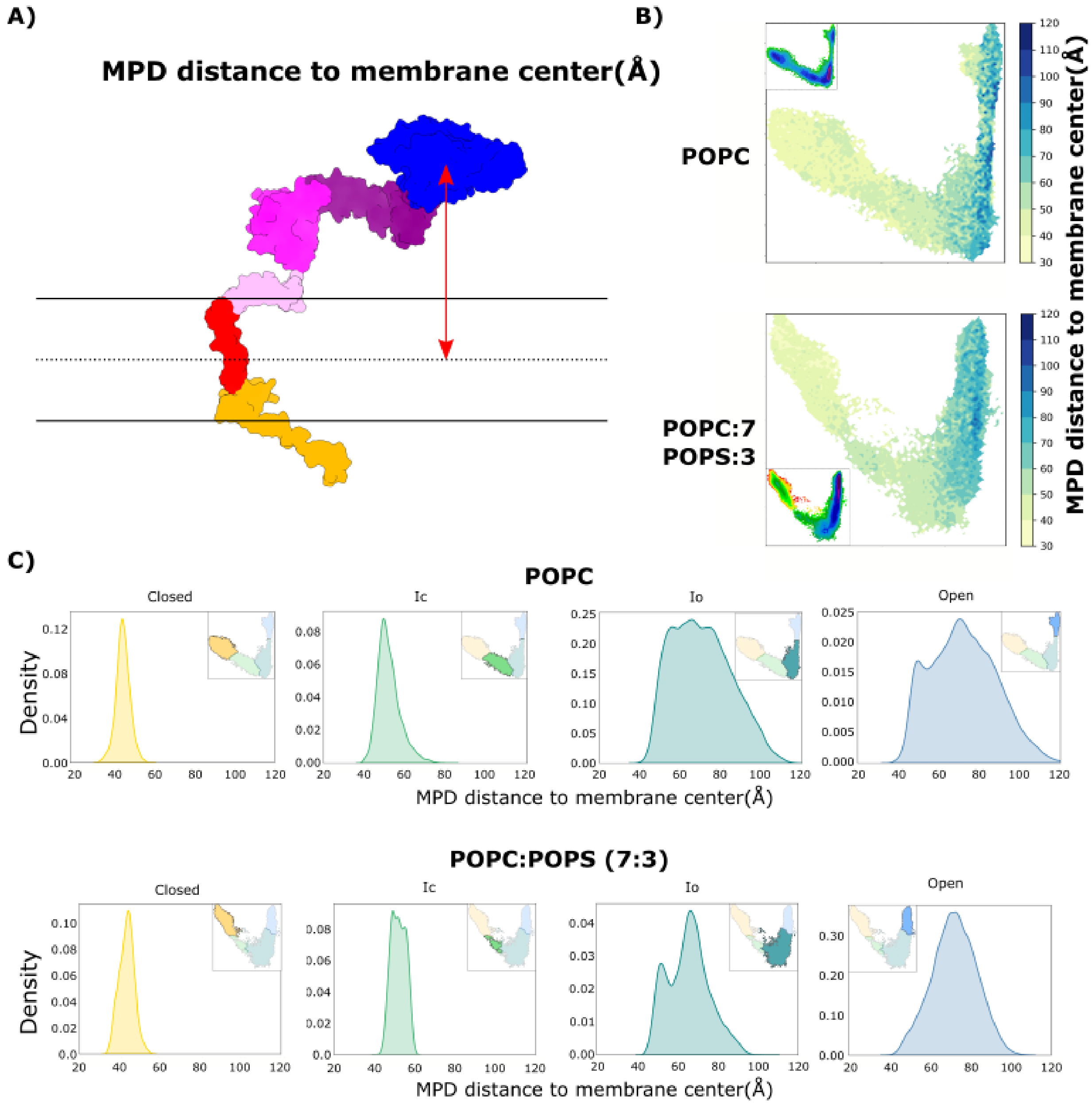


**Supplementary Figure S4. The MpD dissociates from membrane lipids early in ADAM10 opening in the absence or presence of POPS. A)** Representation of the distance between the MpD and the center of the membrane **B)** Projection of this distance over free energy maps in POPC and POPC-POPS simulations. **C)** Distribution of this distance in each macro state obtained in both conditions.

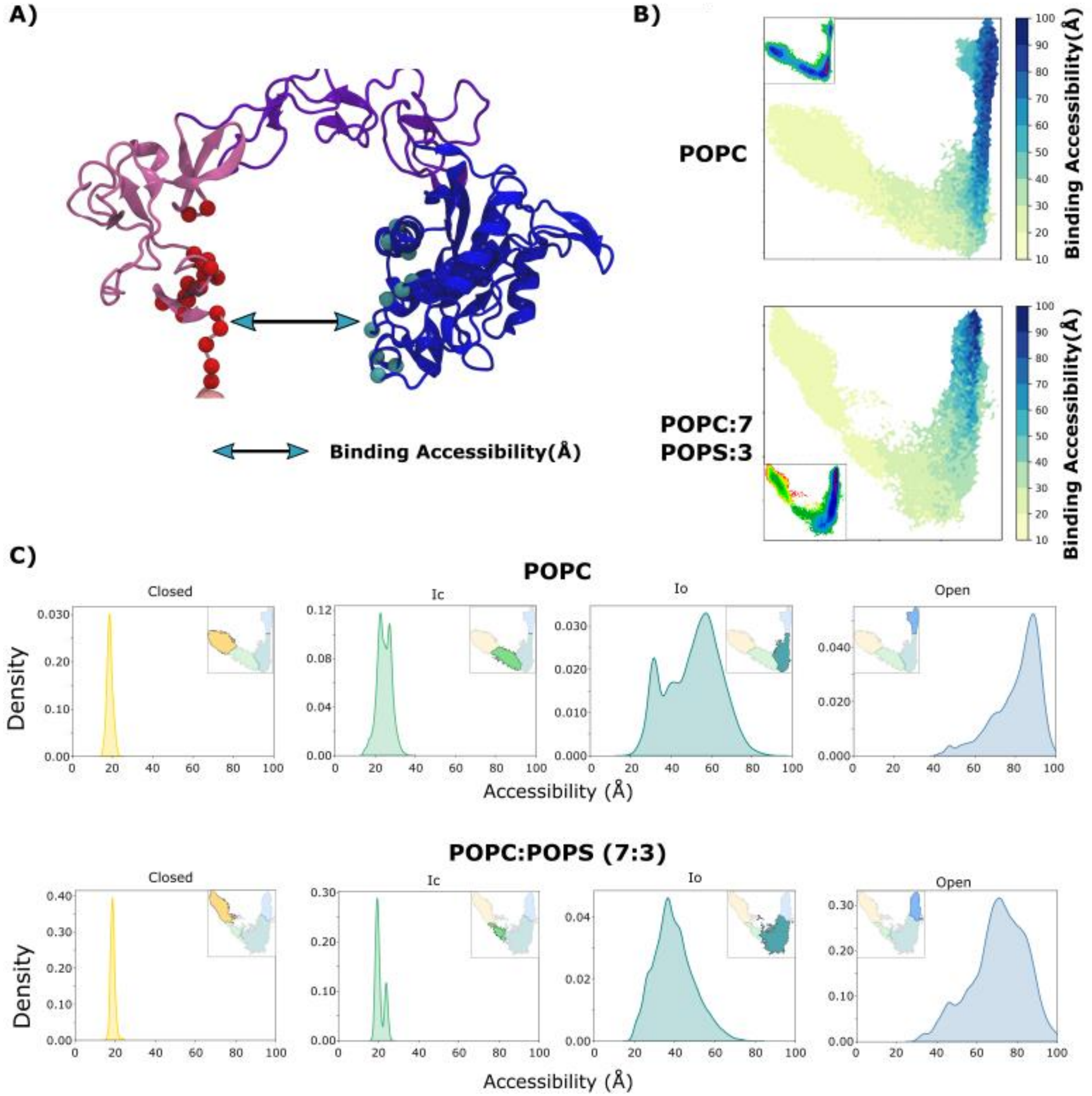


**Supplementary Figure S5. Structural accessibility of ADAM10 to Tspan binding. A)** Representation of the distance defining the accessibility to Tspan **B)** Projection of this distance over free energy maps in POPC and POPC-POPS simulations. **C)** Distribution of this distance in each macro state obtained in both conditions.

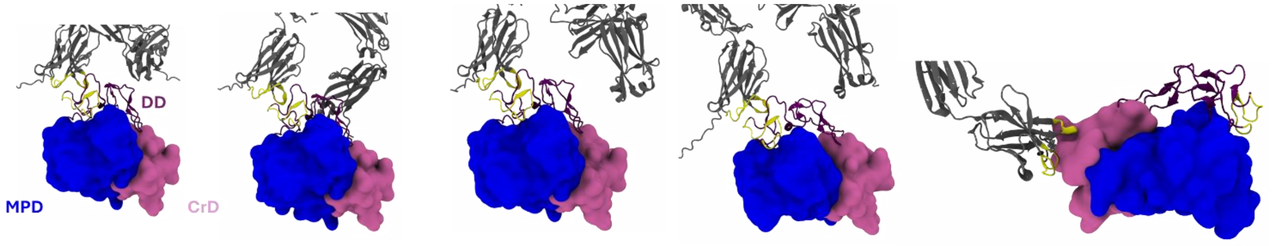


**Supplementary Figure S6: Outputs produced by AF3 when predicting antibody binding to ADAM10.** In 4 of 5 structures generated with AF3, residues loops 48-54, 71-77 and 118-123 from the Fab (in Yellow) interact with the residues 470-498 (DD, in yellow) of ADAM10. In the last predicted model, we observed an interaction between loops 48-54, 71-77 and 118- 123 from the Fab and residues 576-617 (CrD) from ADAM10.

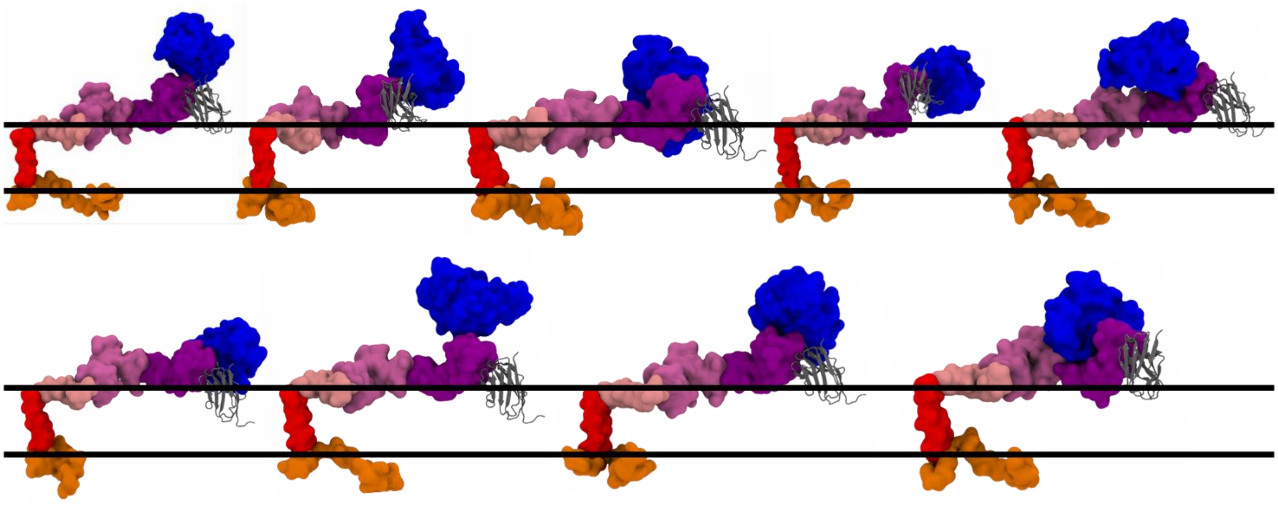

**Supplementary Figure S7: Projection of antibody binding to ADAM10 in most represented state of POPC-POPS condition.** The structure were extracted from the most populated state of POPC:POPS(7:3) simulation using the "sample_by_distributions" routine of pyEmma. The $Fab_{cryo}$ structure was then superimposed with its corresponding binding site on ADAM10 in each of these structures. For clarity, lipids are represented with black lines.

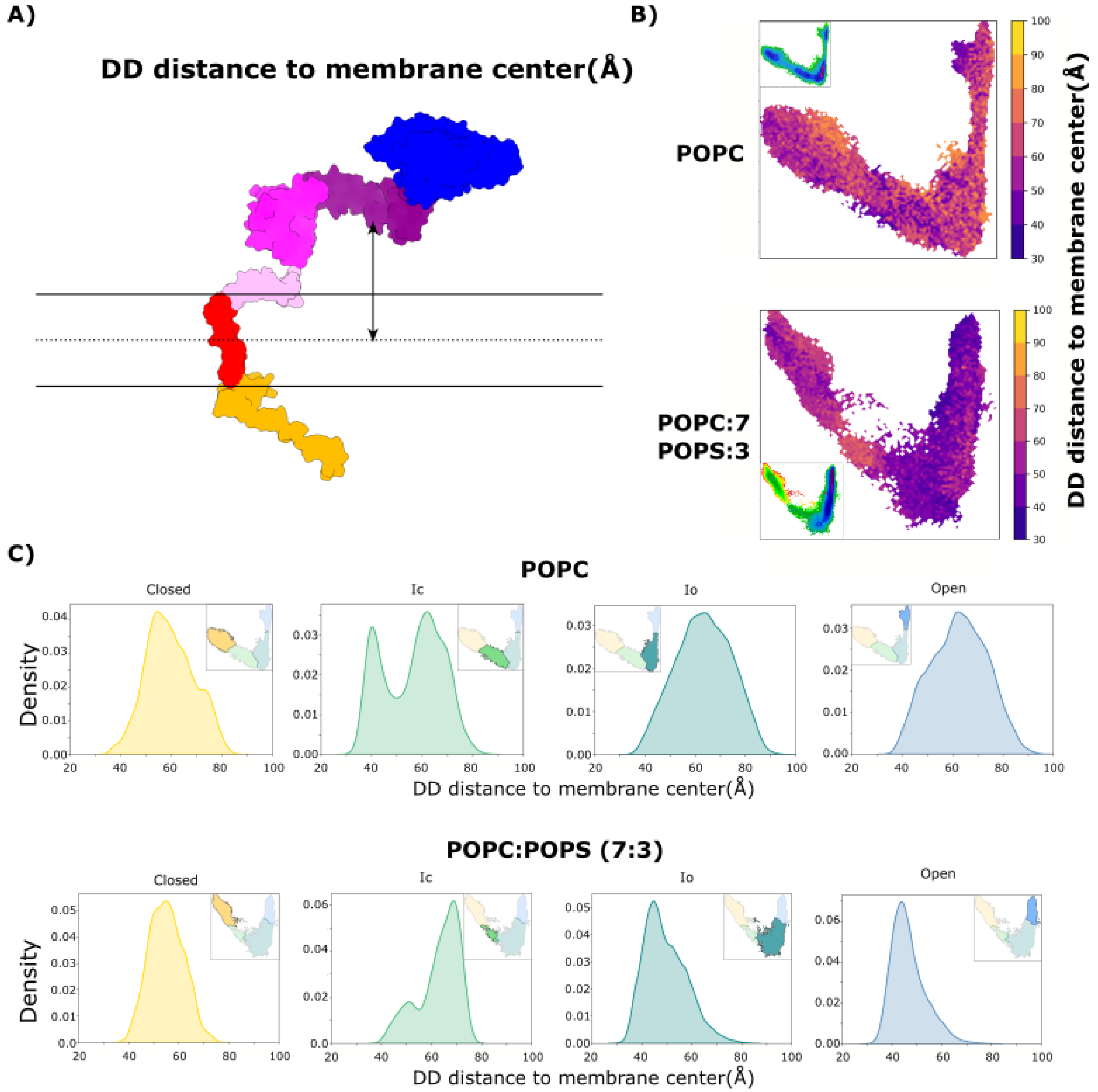


**Supplementary Figure S8. The DD stays closer from membrane lipids in presence of POPS. A)** Representation of the distance between the DD and the center of the membrane **B)** Projection of this distance over free energy maps in POPC and POPC-POPS simulations. **C)** Distribution of this distance in each macro state obtained in both conditions.

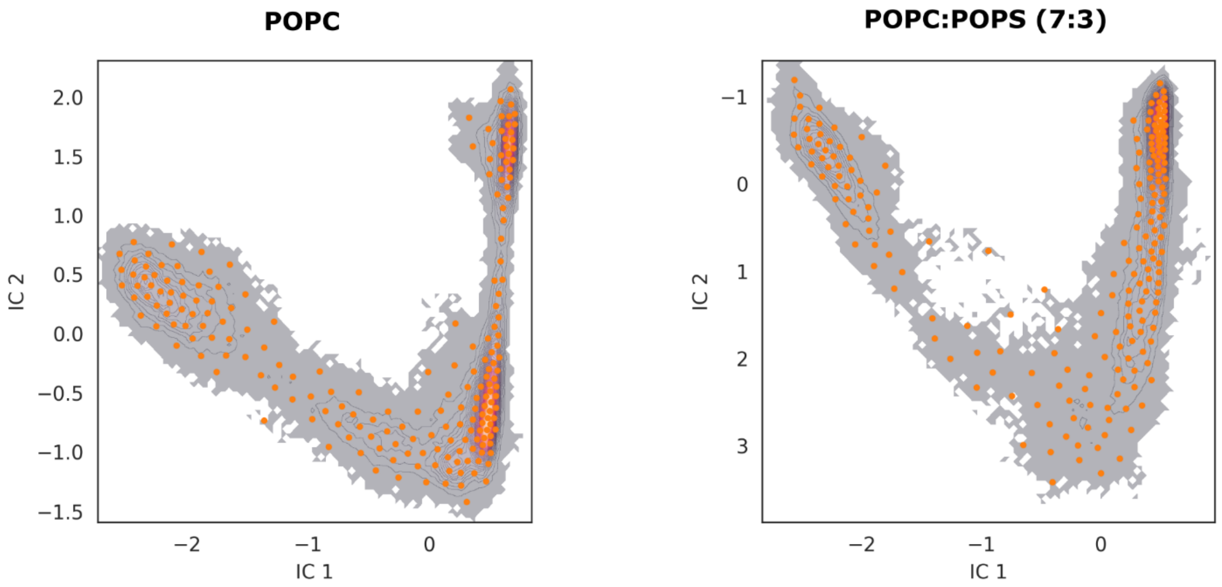


**Supplementary Figure S9. Distribution of cluster center obtained after k-means clustering.** 200 microstates were assigned in both POPC and POPC:POPS(7:3) conditions.

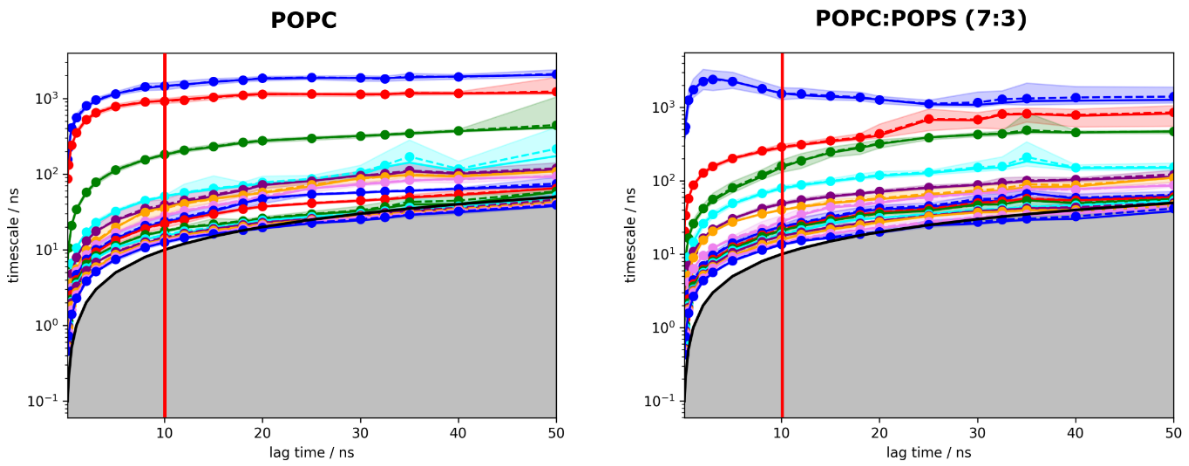


**Supplementary Figure S10. Assessment of convergence of the MSMs with implied timescales.** Implied timescales with MSMs estimated with different lag times for POPC and POPC:POPS(7:3) systems. Red lines correspond to the chosen lag times for analyses.

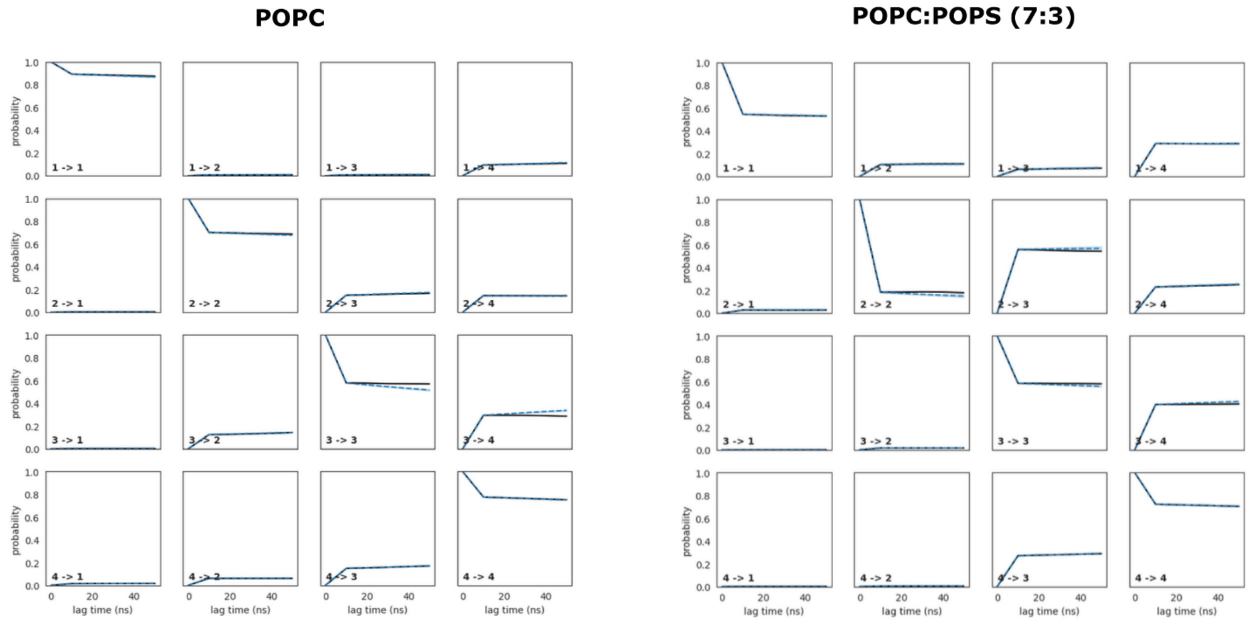


**Supplementary Figure S11. Assessment of convergence of the POPC and POPC:POPS(7:3) MSM with Chapman Kolmogorov tests.** Chapman-Kolmogorov tests with four states for both systems.
